# Fate mapping of peripherally derived macrophages reveals a long-lasting engrafted population that maintains a distinct transcriptomic profile for up to 8 months after Traumatic Brain Injury

**DOI:** 10.1101/2024.11.18.624186

**Authors:** Maria Serena Paladini, Benjamin A. Yang, Xi Feng, Elma S. Frias, Karen Krukowski, Rene Sit, Maurizio Morri, Wendy Lam, Valentina Pedoia, Stefka Tyanova, Susanna Rosi

## Abstract

Traumatic Brain Injury (TBI) is one of the most established environmental risk factors for the development of dementia and long term neurological deficits representing a critical health problem for our society. It is well-established that TBI-induced neuroinflammation contributes to the long-lasting cognitive deficits and engages brain-resident macrophages (microglia) as well as monocytes-derived macrophages (MDMs) recruited from the periphery. While numerous studies have characterized microglia response to TBI, and the critical role of early infiltrated MDMs in the development of cognitive dysfunctions, the fate of MDMs in TBI remains unknown. Microglia and MDMs have distinct embryological origins and it is unclear if MDMs can fully transition to microglia after infiltrating the brain. This gap in knowledge is due to the fact that after brain engraftment, MDMs stop expressing their signature markers, thus making discrimination from resident microglia cells elusive. Here, for the first time, we longitudinally trace the fate of MDMs by taking advantage of two complementary yet distinct fate mapping mouse lines, CCR2-creER^T2^ and Ms4a3-cre, where inflammatory monocytes are permanently labeled even after *in situ* reprogramming. We demonstrated that early infiltrated MDMs persist in the brain for up to 8 months after TBI in adult female and male mice. Notably, MDMs retain their phagocytic activity while remaining transcriptomically distinct from microglia, and show a signature associated with aging and disease. Our data significantly advance the understanding of long-lasting MDMs and provide critical knowledge for developing more targeted therapeutic interventions for myeloid cells.

## 1. Introduction

Traumatic brain injury (TBI) represents a global health challenge, being a major cause of long-term physical, cognitive and emotional disabilities with no treatment available to date ^1,2,3^. TBI has a complex and multimodal pathophysiology characterized by the primary injury, resulting from the initial mechanical insult, and the secondary injury, which encompasses tissue and cell damage that can last from hours to years ^4,5^. Neuroinflammation is a central feature of TBI secondary injury that contributes to long-lasting neurologic deficits in patients ^6,7,8^. Key players in TBI-related neuroinflammation are brain-resident macrophages (further referred to as microglia) and infiltrating immune cells from the peripheral immune system, including monocytes and monocytes-derived macrophages (MDMs). Blood circulating monocytes originate from hematopoietic stem cell progenitors in the bone marrow and can be divided in two major subsets in mice: “classical” Ly6C^high^-Ccr2^+^ monocytes which can infiltrate the inflamed tissues and “non-classical” Ly6C^low^-Ccr2^-^ blood-patrolling monocytes ^9,10,11^. Ly6C^high^-Ccr2^+^ monocytes are thought to be recruited at the site of inflammation, extravasate and differentiate into macrophages (MDMs) after engrafting the tissue. The influx of monocytes in the brain is mainly driven by chemokines, a family of chemoattractant cytokines, that are released by resident cells including astrocytes and microglia ^12^, and induce directional migration of peripheral immune cells into the injury site. The key chemotactic role of CC ligand-2 (CCL2, formerly MCP-1) and its receptor Ccr2, has been well-established in the context of TBI. Increased levels of CCL2 have been found acutely in the cerebrospinal fluid ^13,14^ and chronically in the serum ^15^ of patients after TBI. CCL2 increases were also observed in the serum ^16^ and in the lesioned cortex ^13,16,17^ of TBI mice. Moreover, blocking Ccr2 either genetically by using Ccr2^-/-^ mice ^13,18,19,20,21^ or pharmacologically by using Ccr2 antagonists ^16,22,23^ dampens monocyte infiltration and prevents TBI-induced cognitive deficits in mice. While these previous studies demonstrated a causal role for monocytes in the damaging effects of neuroinflammation following TBI and the development of cognitive deficits, nothing is known on the long-term fate of engrafted MDMs.

The longitudinal trajectories of MDMs and their contribution to tissue-resident macrophages remains a subject of ongoing debate, as well as their different ontogeny and tissue-specific function in homeostatic condition and during disease ^24^. In organs such as the heart, pancreas and gut, circulating monocytes can give rise to and maintain the resident tissue macrophages population at steady state. In the brain, microglia originate from the yolk sac and continue to self-maintain with minimal input from infiltrating peripheral cells ^25,26,27^. However, during disease, specific chemokine signaling promotes the recruitment of peripheral monocytes into the brain parenchyma and their differentiation into macrophages. Some studies suggested that monocytes invading the brain do not contribute to the resident microglia pool ^28, 29, 30^, while other groups described that monocytes transition to microglia ^31,32^. A key limitation of these studies is the lack of reliable tools for precise cell tracing post-engraftment, which leaves the true ontogeny and identity of infiltrating monocytes and macrophages unclear. Recently developed fate mapping mouse models that enable accurate lineage tracing of monocytes are useful tools to unravel the role of MDMs in diseases and homeostasis, especially in comparison to microglia ^33^. Two leading models for the tracing of monocytes are driven by *Ccr2* and *Ms4a3* promoters (specifically expressed by monocyte-committed bone marrow progenitors) ^34–36^. Here we used Ccr2-creER^T2^ and Ms4a3-cre mice to unambiguously trace blood-recruited MDMs 7, 30 days and 8 months after TBI. We determined the spatial localization, transcriptomic signature and functionality of MDMs focusing on how these features change at subacute and chronic time points and how they differ from resident microglia. Moreover, we identified the shared core transcriptomic signature of infiltrated macrophages across different experimental conditions that will be crucial for the development of therapeutic strategies aimed to specifically target these cells.

## 2. Methods

### 2.1. Animals

All experiments were conducted in accordance with National Institutes of Health (NIH) Guide for the Care and Use of Laboratory Animals and approved by the Institutional Animal Care and Use Committee (IACUC) of both the University of California, San Francisco (AN184326) and from Altos Labs, Inc (EB22-101-100) to ensure compliance with ethical standards and the humane treatment of animals.

Ccr2-creER^T2^-mKate2 mice were obtained from the University of Zurich (UZH). Ms4a3-cre were obtained from the Singapore Immunology Network (SIgN), A*STAR. Ai14D reporter mice were purchased from Jackson laboratory (#007914). Ccr2-creER^T2^-mKate2 and Ms4a3-cre mice were crossed with Ai14D mice to obtain Ccr2-creER^T2^::Ai14D and Ms4a3-cre::Ai14D mice. Female and male mice from both genotypes were 11-17 weeks of age at the time of surgeries. Mice were group housed (by sexes and injury state) in environmentally controlled conditions with a reverse light cycle (12:12 h light: dark cycle at 21 ± 1 ◦C; ∼50% humidity) and provided food and water ad libitum.

### 2.2. Tamoxifen treatment and Traumatic Brain Injury surgery

Tamoxifen (Sigma, #T5648) was dissolved in corn oil and administered daily to Ccr2-creER^T2^::Ai14D mice by intraperitoneal injection of 5mg/40g body weight for 5 days^37^. To achieve the desired labeling of inflammatory monocytes (Ccr2+) in the blood, Ccr2-creER^T2^::Ai14D mice were treated with tamoxifen for 3 days before injury, right after injury and the day after, for a total of 5 injections.

Ccr2-creER^T2^::Ai14D and Ms4a3-cre::Ai14D mice were randomly assigned to each TBI or sham surgery group. Animals were anesthetized and maintained at 2–2.5% isoflurane. Controlled Cortical Impact (CCI) surgery was performed as previously described^19,38^. Briefly, mice were secured to a stereotaxic frame with nontraumatic ear bars. A midline incision exposed the skull followed by a ∼3.5-mm diameter craniectomy and removal of part of the skull, using a manual microdrill or a robot microdrill (Neurostar GmbH, customized). The coordinates of the craniectomy were anteroposterior, -2.0 mm and mediolateral, + 2.0mm with respect to bregma. Any animal that experienced excessive bleeding due to disruption of the dura was excluded from the study. After the craniectomy, the removed skull was discarded, and the contusion was induced using a 3 mm convex tip attached to an electromagnetic impactor (Leica). The contusion depth was set to 0.95 mm from dura with a velocity of 4.0 m/s sustained for 300 ms. Following impact, the scalp was sutured. These injury parameters were chosen to target, but not penetrate, the hippocampus. Sham animals underwent a similar procedure but without removal of the skull and no impact. Post-surgery, the mice recovered in a cage on top of a heated pad until they showed normal walking and grooming behavior, and then returned to their home cages. Only animals that fully recovered from the surgical procedures as exhibited by normal behavior, healed sutures, and weight maintenance monitored throughout the duration of the experiments were used.

### 2.3. Radial arm water maze

The radial arm water maze (RAWM) task was used to test spatial learning and memory ^38,39^. Pool dimensions were 118.5 cm in diameter with 8 arms, each 41 cm in length, and an escape platform. The escape platform was slightly submerged below the water level and water was rendered opaque by adding white paint (Crayola, #54–2128–053), so it was not visible to the animals. Visual cues were placed around the room. Two different versions of RAWM were used, a two-days version at 7 days post injury (dpi) and a three-days version at 30 dpi and 8 months after injury. In the first version, mice performed 9 trials on the learning day and 3 trials during the memory probe, 24h after. In the second version, animals ran 6 trials/day during learning and 3 during the memory probe, one week later. On both versions, during both learning and memory days there was a 10 min inter-trial interval. During a trial, animals were placed in a random arm (excluding the arm where the escape platform is located). Animals were allowed 1 min to locate the escape platform. On successfully finding the platform, animals remained there for 10 s before being returned to their warmed, holding cage. On a failed trial, animals were guided to the escape platform and then returned to their holding cage 10 s later. The escape platform location was the same in each experimental cohort, whereas the start arm varied between trials. The number of errors from 3 consequent trials were averaged in one “block”. Data was graphed as the number of non-targeted entries (“number of errors”).

### 2.4. Flow Cytometry

#### Blood

To test efficiency of tamoxifen-induced labeling of peripheral Ccr2+ monocytes, blood was collected from Ccr2-creER^T2^::Ai14D via tail vein puncture. A small nick was made in the tail vein using a scalpel and ∼ 40µl of blood was removed using a pipette and placed in a small tube containing EDTA (Sigma, #E8008). In Ms4a3-cre::Ai14D mice, ∼200µl of blood was extracted by cardiac puncture once completely anesthetized before tissue collection. Red blood cells were lysate using RBC lysis buffer (BioLegend, #420301) and samples were then blocked with CD16/32 Fc block (BD Biosciences #553141) and stained with fluorophore-conjugated antibodies: CD11b-AF700 (BD Pharmingen, #557690), CD11b-BV421 (BD Pharmingen, #562605), CD45-FITC (BD Pharmingen, #553080), CD45-APC-Cy7 (BioLegend, #304014), Ly6C-V450 (BD Pharmingen, #560594), Ly6C-APC (BD Pharmingen, #560600) Ly6G-PE-Cy7 (BD Pharmingen,# 560601) and Ly6G-BV711 (BD Pharmingen, #563979). Cells were then washed in FACS buffer (1×DPBS with 0.5% BSA fraction V and 2% FBS) and used for analyses. Data were collected on BD FACSymphony™ A3 Cell Analyzer and on FACSAria^TM^ (III and Fusion) cell sorters (BD Biosciences, V8.0.1), and analyzed with FlowJo software (FlowJo, LLC).

#### Brain

Mice were euthanized and perfused with cold PBS. Brains were immediately removed and dissociated using a Neural Tissue Dissociation kit (P) (Miltenyi Biotec, #130-092-628). Cells were then resuspended in 30% Percoll solution diluted in RPMI medium, and centrifuged at 800 g for 20 min at 4 °C. Cell pellets were washed with FACS buffer (1×DPBS with 0.5% BSA fraction V and 2% FBS or Rockland, #MB-086-0500), blocked with mouse CD16/32 Fc block (BD Biosciences, #553141) and then stained with fluorophore-conjugated antibodies: CD11b-AF700 (BD Pharmingen, #557690), CD11b-BV421 (BD Pharmingen, #562605), CD45-FITC (BD Pharmingen, #553080), CD45-APC-Cy7 (BioLegend, #304014), Ly6C-V450 (BD Pharmingen, #560594), Ly6C-APC (BD Pharmingen, #560600) Ly6G-PE-Cy7 (BD Pharmingen,# 560601) and Ly6G-BV711 (BD Pharmingen, #563979). Cells were then washed in FACS buffer and used for analyses or sorted. Data were collected on BD FACSymphony™ A3 Cell Analyzer and on BD FACSAria^TM^ (III and Fusion) cell sorters (BD Biosciences, V8.0.1), and analyzed with FlowJo software (FlowJo, LLC).

### 2.5. Immunofluorescence

For immunohistochemistry analysis, animals were lethally overdosed, followed by PBS perfusion. Brains were fixed in ice-cold 4% paraformaldehyde, pH 7.5 (PFA, Sigma Aldrich, St. Louis, MO, 441244) overnight, cryo-protected in 15% and 30% sucrose (Fisher Scientific) and sectioned into 20 μm coronal slices using a Leica cryostat (Leica Microsystems). Slides were then placed in 100% methanol at -20C for 10 minutes, blocked in BSA 5% in TBS-Tween and then stained with antibodies (488-conjugated rabbit anti-Iba1, Cell Signaling, #20825, 1:100 and anti-P2yr12, Anaspec, #AS-55043A, 1:400) overnight at 4C and then 2h at room temperature with secondary antibodies. Nuclei were stained with DAPI. Tissues were fixed using ProLong Gold (Invitrogen, #P36930) and a standard slide cover sealed with nail polish.

Pericontusional regions (top quarter of a coronal brain section) were analyzed by acquiring z-stacked tiled images on a Axioscan 7 slide scanner (Zeiss) using a 20x lens. For each mouse, cavitation images were acquired at 3 different coordinates from rostral to caudal: approximately Bregma -1.34mm (“Anterior”), -1.81mm (“Central”) and -2.54mm (“Posterior”).

Imaris software (Imaris 10.2.0) was used for machine learning-based surface rendering of Iba1, P2yr12, DAPI staining and tdTomato signal. A tdTomato/DAPI overlapped volume ratio of 0.5 was used as threshold criteria to include cells in the analysis. Cell volume data were exported and analyzed.

### 2.6. In vivo phagocytic assay

To test phagocytosis capacity of infiltrated macrophages, 2μl of 2μm latex beads (Sigma Aldrich, #L0280) diluted 1:10 in saline were injected into the right hippocampus (ipsilateral to the CCI injury) using the following coordinate: bregma, AP − 1.6 mm, ML + 1.6 mm and DV − 2mm. Mice were euthanized three days post injection and phagocytosis was evaluated by immunofluorescent staining (colocalization of blue beads in tdTomato+ (Ccr2+) cells). Nuclei were stained with H3K9me3 antibody (abcam, #ab176916) and visualized in green, as the blue channel was used by the latex beads. 40x Z-stack tiled images of the injection site (top half of the injured hemibrain) were acquired on a Axioscan 7 slide scanner (Zeiss). 2/4 images were acquired for each animal. One representative image was acquired using a LSM980-Airyscan2 (Zeiss) laser scanning confocal. Maximum Intensity Projection and normalization to the 1st and 99th percentile was performed for each channel of z-stack. tdTomato+ macrophages and blue beads were automatically segmented using the 2-class K-means algorithm implemented in matlab. Circularity index was computed as (4*pi*Area/Perimeter^2)*(1 - 0.5/r)^2 where r = Perimeter/(2*pi) + 0.5, for each segmented connected component in the tdTomato channel. Blobs with circularity index <0.7 were excluded by the analysis as well as cells with cross sectional area outside the range of 5-20 μm. Intersection between each remaining Ccr2+ cell and the blue beads segmentation mask was computed and each cell with non zero intersection was counted as positive in blue beads and tdTomato+ cell colocalization. Data are graphed as % of tdTomato+ macrophages engulfing one or more beads (number of macrophages that colocalize with beads in the acquired image/ total number of macrophages in the image*100).

### 2.7. Bulk RNA sequencing

#### Library Preparation

CD11b+ CD45+ tdTomato+ cells (MDMs) were sorted from whole brains of Ccr2-creER^T2^::Ai14D mice at one week, one month and eight months after injury and from Ms4a3-cre::Ai14D mice at one week and one month after injury. For the first two time points (7 and 30 days), samples were processed as follows. cDNAs were generated using the SMART-Seq v4 Ultra Low input RNA kit for sequencing (Takara bio, 634891), according to the vendor protocol with 14 cycles of cDNA amplification. At the last step, the cDNA was eluted in 30ul of RNase/DNase free water. Subsequently 30ul of cDNA was used as a starting material with Illumina DNA Prep library kit (Illumina, 20018705) according to the vendor protocol, with final library PCR amplification of 10 cycles. After library completion, individual libraries were pooled equally by volume, and quantified on Fragment Analyzer (Agilent, DNF-474). Quantified library pool was diluted to 1nM and sequenced on MiniSeq (Illumina, FC-420-1001) to check for quality of reads. Finally, individual libraries were normalized according to MiniSeq output reads, specifically by % protein coding genes and were sequenced on two lanes of HiSeq4000 SE50 for a total of ∼30M reads per sample.

For the 8 months time point (Ccr2-creER^T2^::Ai14D mice) and the Ms4a3-cre::Ai14D mice, cDNAs were generated using the SMART-Seq® mRNA LP (with UMIs) kit for sequencing (Takara bio,634765), according to the vendor protocol. cDNA amplification cycle was adjusted to 19 cycles. PCR-amplified cDNA was purified by immobilization on NucleoMag NGS Clean-up and Size Select beads. The beads were then washed with 80% ethanol, and cDNA is eluted with ∼15ul of Elution Buffer. Subsequently, 8 ul of cDNA 0.125 ng/μL was used as a starting material for library preparation using the the Unique Dual Index (UDI) kits (Takara bio, 634752-5) according to the vendor protocol, with final library PCR amplification of 14 cycles. PCR-amplified library was purified individually by immobilization on NucleoMag NGS Clean-up and Size Select beads. The beads were then washed with 80% ethanol and the library eluted with nuclease-free water.

Quality control and quantification was performed using the Qubit dsDNA HS Assay Kit (Thermofisher, 32854) and TapeStation D5000 Reagents (Agilent, 5067-5589) and Screentape (Agilent, 5067-5588). Samples were normalized and pooled individually for sequencing. An additional double-sided SPRI selection was performed on the pool prior to sequencing. Libraries were sequenced using the NextSeq 2000 P3, 300 cycles kit (Illumina, 20040561).

#### Pre-processing and QC

RNA-seq FASTQ files were trimmed, aligned to the mouse reference genome (GRCm39, v108), and quantified using the nf-core/rnaseq pipeline with the star_salmon option (v3.12.0)^40^. Gene-level length-scaled counts were used for all downstream analyses. In the 8-month Ccr2 and Ms4a3 datasets, UMIs were trimmed from all reads before alignment and quantification, and latent batch effects were identified and removed using Surrogate Variable Analysis (SVA)^41^. GC-content correction offsets were calculated for the Ms4a3 datasets using the cqn package^42^.

#### Differential gene expression

Differentially expressed genes (DEG) between conditions at each time point were identified within each dataset using the quasi-likelihood framework implemented in edgeR. Raw length-scaled counts were filtered to retain genes with a minimum of 10 counts within biological groups and trimmed-mean of M-value (TMM) normalization factors were calculated. Empirical Bayes estimates of the quasi-likelihood dispersions were robustified against outlier genes. Thresholds of adjusted p-value < 0.05 and absolute log-fold-change > 0 were used to identify differentially expressed genes. Differential expression testing was not performed between datasets (Ccr2 7dpi & 30dpi, Ccr2 8mos, Ms4a3) due to batch confounding.

#### Single-sample scoring (singscore)

To compare groups across batch-confounded datasets, the filtered counts tables from all datasets were merged using the intersection of expressed genes. Genes were then ranked by expression within each sample and independently scored for enrichment against published transcriptomic signatures and gene sets from several databases (i.e. Reactome^43^, ImmuneSigDB and Cell Type Signature Signatures from MSigDB^44–46^) using the singscore package^47^. The Reactome terms were downloaded from https://www.reactome.org. The ImmuneSigDB and Cell Type Signatures were collected from the msigdbr package. Enrichment scores were then compared between groups and datasets. The DIMs and DAMs signatures were collected from Supplementary Table 1 of Silvin et al., 2022^48^. The SenMayo signature was collected from Supplementary Data 1 of Saul et al., 2022^49^. The Microglia Aging and Accelerated (Accel.) Microglia Aging signatures were collected from Holtman et al., 2015^50^ as stored in the 2021 HDSigDB database at https://www.maayanlab.cloud.

#### Sparse partial least squares discriminant analysis (sPLS-DA)

First, PLS-DA was performed using the mixOmics package^51^ for singscore scores from each database (i.e. Reactome, MSigDB cell type signatures, ImmuneSigDB) using at least 10 components. Model performance was assessed using M-fold cross-validation with the “perf” command with 50 repetitions. 5 folds were used for the Ccr2 datasets and 3 folds were used for combined analysis of the Ccr2 and Ms4a3 datasets. Next, sparse PLS-DA (sPLS-DA) was performed using one additional component than the optimal number of components identified from PLS-DA. Cross-fold validation was performed to determine the optimal number of components and important features per component with the same number of folds and repetitions. The resulting number of components and features were used to train the final sPLS-DA model. 3 components were selected for the Ccr2 analysis and 9 components were selected for the combined analysis of Ccr2 and Ms4a3 with the cell type signatures database. The selected features for each component were visualized using heatmaps of singscore z-scores for all terms with non-zero contributions to any component.

#### Overlap with core MDM signature

The core MDM signature from Du et al., 2024 was overlapped with differentially expressed genes identified in several datasets. “Ms4a3 TBI Mac vs Mic” refers to the union of genes that are upregulated in TBI macrophages relative to TBI macrophages in the Ms4a3 model across 7dpi and 30dpi. “Feng et al., 2021” refers to genes that are upregulated in brain-engrafted macrophages collected from irradiated mice treated with PLX versus microglia collected from control mice without PLX. “Ccr2 8mos TBI Mac vs Mic” refers to genes upregulated in 8-month TBI macrophages versus 8-month TBI microglia. UpSet plots were also created using the ComplexHeatmap package.

The 29 overlapping MDM signature genes were annotated to Gene Ontology (BP) terms using the goana function in limma. The universe was set as genes that were commonly expressed across all of the examined datasets except those analyzed in Du et al., 2024 (15,985 genes).

#### Heatmaps and volcano plots

Heatmaps were created using the ComplexHeatmap package^52^ with z-scores of logCPM values or singscore scores. Volcano plots were created using the EnhancedVolcano package.

### 2.8. Statistical Analyses

All data were analyzed with GraphPad Prism 9 statistical software. Cognitive performance in the RAWM maze was analyzed as a two-way repeated measure (RM) analysis of variance (ANOVA) followed when appropriate by Šidák multiple comparisons test. Differences in phagocytosis were analyzed using Unpaired T-test. Cell volume, data across time points were analyzed using one-way ANOVA followed by Tuckey post hoc test or Unpaired T-test.

Group outliers were determined (ROUT method, Q=1%) and excluded from analysis. P values below 0.05 were considered significant. Individual animal scores represented by dots, lines depict data mean and SEM. Individual statistical analysis is denoted in the figure legends.

For statistical analysis on the transcriptomic data, refer to the corresponding Methods section.

#### Software and Algorithms

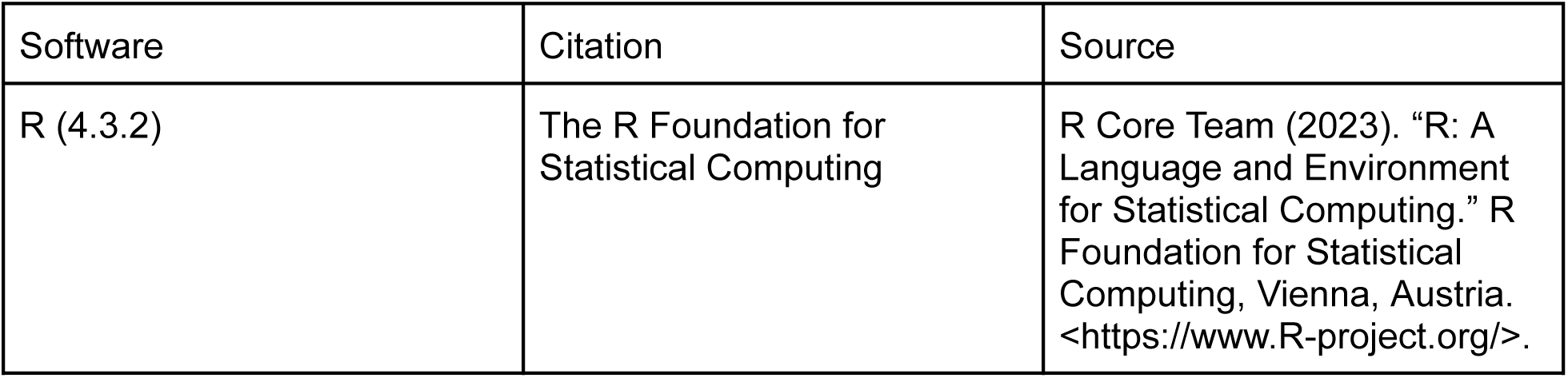
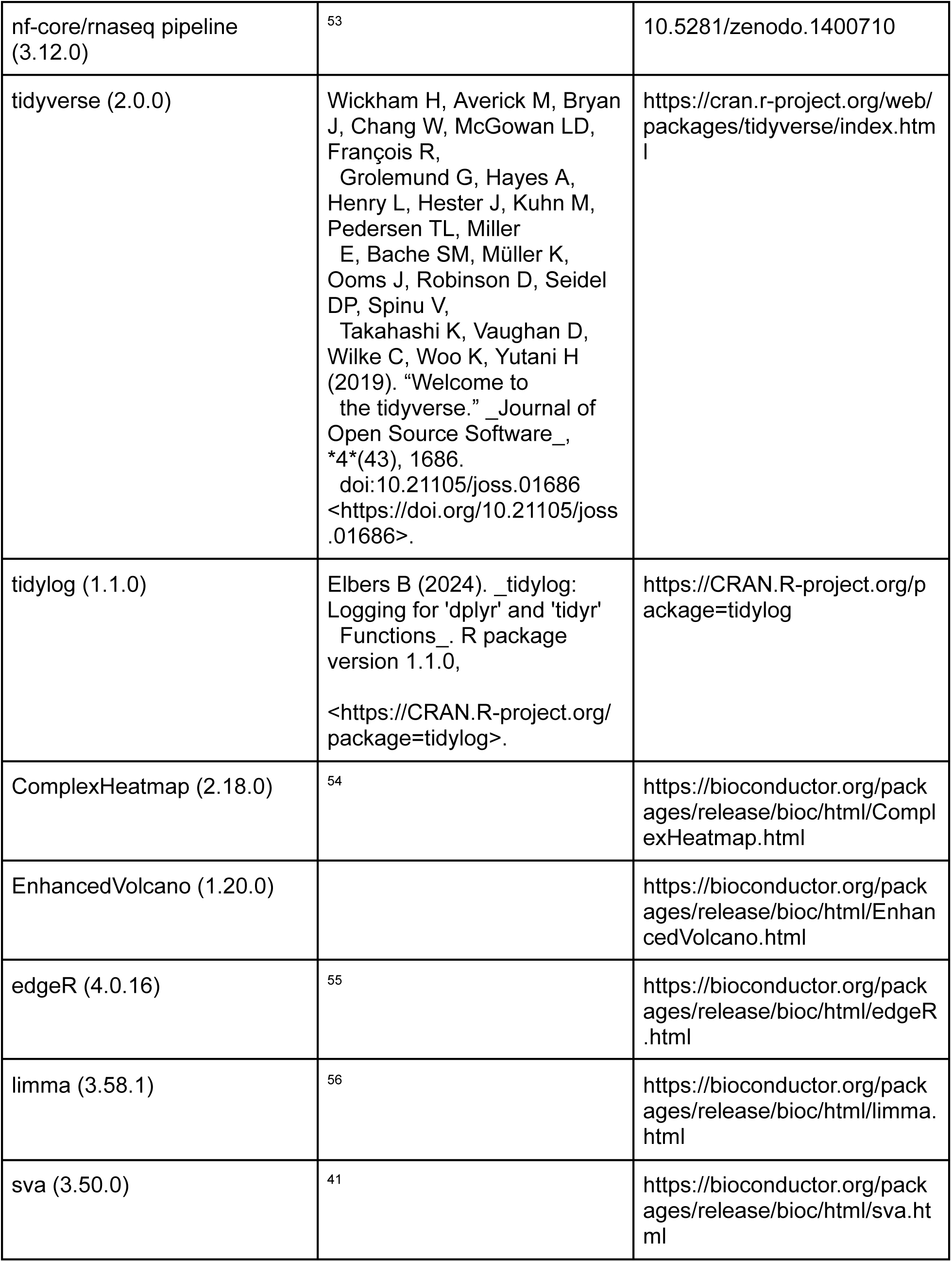
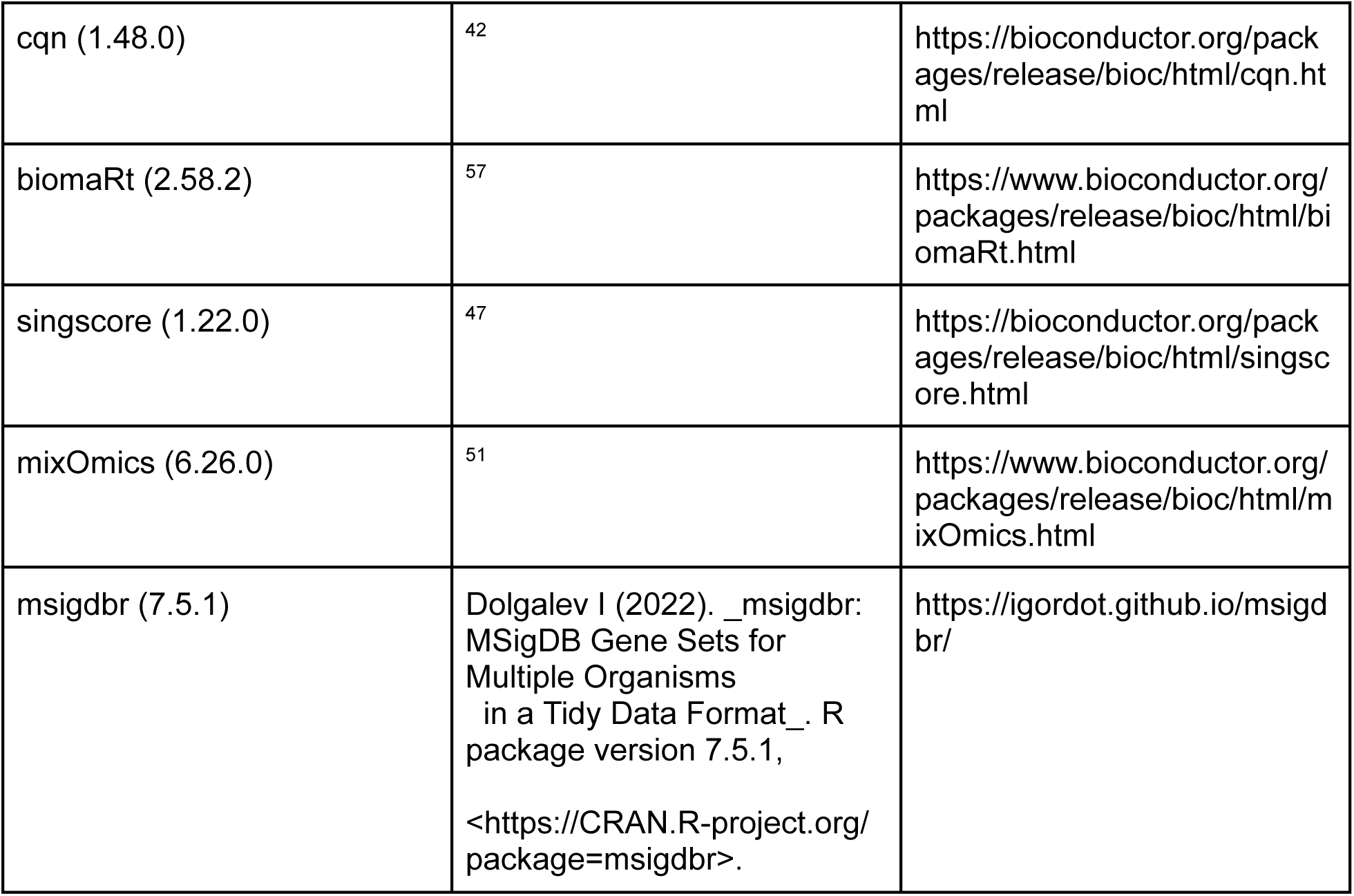

## 3. Results

### 3.1 MDMs infiltrate the injured brain acutely after TBI and persist for up to 8 months

To investigate the fate of MDMs after TBI, Ccr2-creER^T2^ mice were crossed with Ai14D mice to allow for inducible, specific and temporally-controlled labeling of Ly6C^high^-Ccr2^+^ monocytes upon tamoxifen administration ^35,36^. Given the fast turnover of Ly6C^high^ monocytes^58^, we first tested the labeling efficiency of Ly6C^high^ monocytes in the blood after 5 daily tamoxifen injections by flow cytometry (Figure 1A). Starting from 3 doses of tamoxifen and up to two days after stopping treatment (“2 days off”), the % of Ly6C^high^ monocytes that were tdTomato+ (labeling efficiency) was higher than 80% (Figure 1B). Of note, due to the rapid turnover of Ly6C^high^ monocytes, the % of labeled Ly6C^hi^ monocytes decreased to around 25% one week after the last tamoxifen injection, and to almost zero after two weeks (Figure 1B). These results demonstrate that the Ccr2-creERT2::Ai14D system enables precise, temporally-controlled labeling, allowing for the accurate tracing of specific monocyte populations at distinct time points.

**Figure 1.**
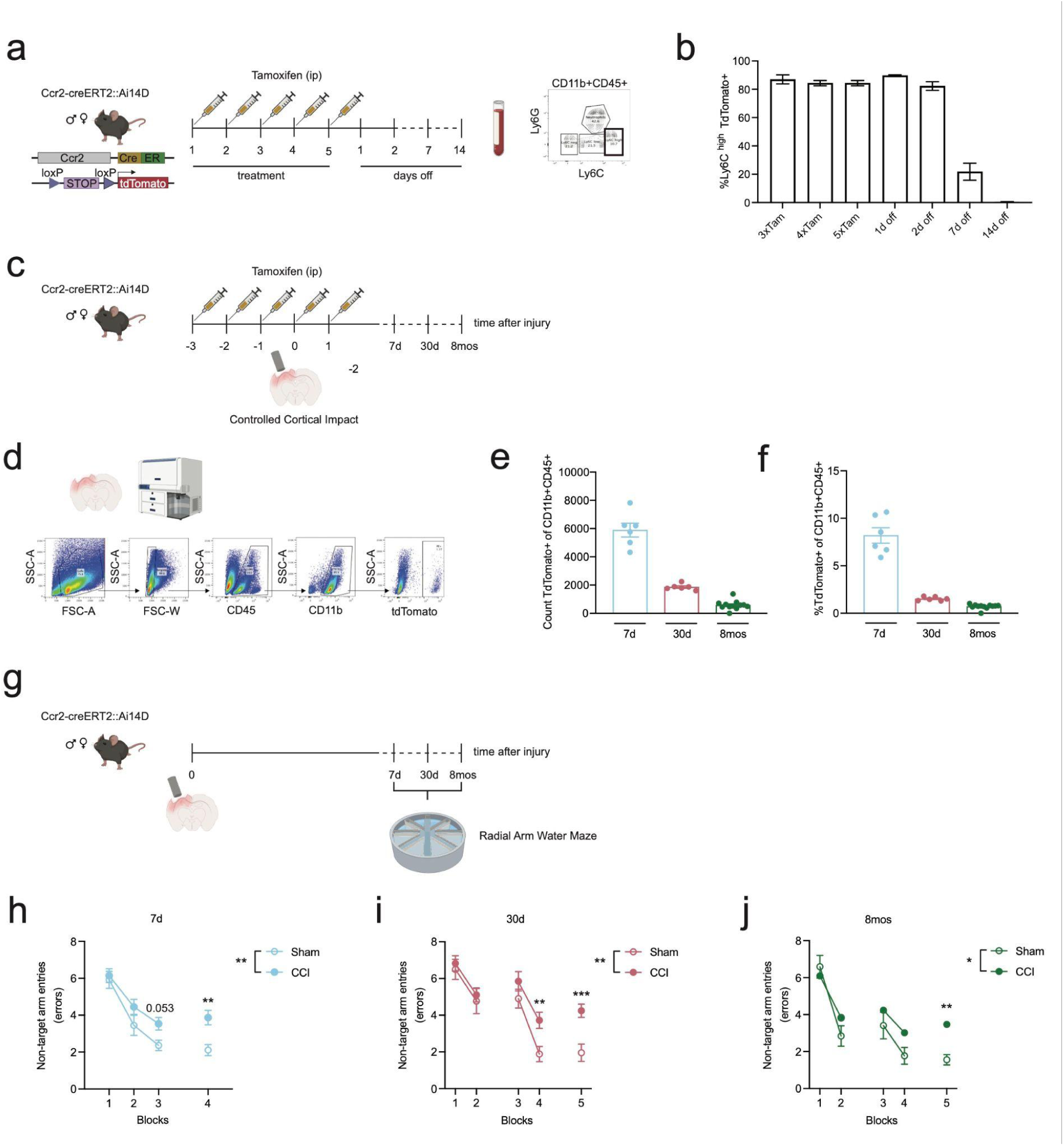
MDMs accumulate and persist in the mouse brain for several months after TBI, paralleled by long-lasting cognitive deficits. (a) Experimental design - tamoxifen-induced labeling of Ly6C^hi^-Ccr2+ blood monocyte in Ccr2-creER^T2^::Ai14D mice. Blood from the tail vein was collected at different time point and analyzed by flow cytometry (b) Percentage of blood Ly6C^hi^ monocytes that were efficiently labeled (tdTomato+) in male and female Ccr2-creER^T2^::Ai14D mice (c) Experimental design - female and male Ccr2-creER^T2^::Ai14D mice received a total of 5 doses of tamoxifen and were injured using the Controlled Cortical Impact model of TBI on the fourth day of tamoxifen treatment. (d) Gating strategy for quantifying Ccr2+ (tdTomato+) cells from the injured brain of Ccr2-creER^T2^::Ai14D mice at 7 days, 30 days and 8 months after TBI (e) Average cell number of Ccr2+ (tdTomato+) cells in the CD11b+CD45+ population (f) Percentage of Ccr2+ (tdTomato+) cells in the CD11b+CD45+ population. (g) Experimental design - Ccr2-creER^T2^::Ai14D mice were tested in the Radial Arm Water Maze (RAWM) task to detect TBI-induced cognitive deficits at different time point (h - j) Injured Ccr2-creER^T2^::Ai14D mice made significantly more errors when performing the RAWM task compared to sham (Two-way RM ANOVA revealed TBI effect *p* = 0.0039 and time effect *p* < 0.0001 at 7 days; TBI effect *p* = 0.0041 and time effect *p* < 0.0001 at 30 days and TBI effect *p* = 0.0220 and time effect *p* < 0.0001 at 8 months after injury). Statistical differences from multiple comparison tests are denoted in the graphs. Data are expressed as the mean of the examined variable ± SEM (B, of *n* = 6/21 mice/group). \**p*<0.05, \*\**p*<0.01, \*\*\**p*<0.001 (Two way RM ANOVA with Šidák multiple comparisons test).

We have previously shown that the accumulation of MDMs in the ipsilateral hemisphere starts at 12 hours and peaks at 24 hours after TBI^16^. To confidently capture and follow solely the initial infiltration window of Ly6C^high^ monocytes, the Controlled Cortical Impact (CCI) model was used to induce TBI in mice that received three out of the five total doses of tamoxifen (Figure 1C). At this time point (0 dpi, 3 tamoxifen doses), the labeling efficiency was between 80 - 85%, with no sex differences observed (Supplementary Figure 1A). Of note, off-target labeling of neutrophils (Ly6C+Ly6G+) and patrolling monocytes (Ly6C^low^) was negligible (Supplementary Figure 1B). We next evaluated TBI-induced infiltration and engraftment of labeled Ccr2+ cells by flow cytometry on whole brain samples at different time points (Figure 1D). As expected we measured labeled MDMs in the injured brain after 7 days (average count: 5890.5 ± 490 cells, 8.2 ± 0.81% of the CD11b+CD45+ whole brain population), in line with our previous work ^16^. Surprisingly, MDMs that infiltrated acutely after TBI were still present after 1 month (average count: 1862 ± 87 cells, 1.52 ± 0.08 % of the CD11b+CD45+ whole brain population) and 8 months (average count: 580.3 ± 92 cells, 0.71 ± 0.07% of the CD11b+CD45+ whole brain population) (Figures 1E and 1F) post TBI. These data provide the first evidence that engrafted MDMs can last in the injured brain for several months following a contusion injury.

### 3.2 TBI-induced cognitive deficits last for up to 8 months in Ccr2-creER^T2^::Ai14D mice

Given the long-lasting presence of early infiltrated MDMs and previous work demonstrating their causal role in the development of TBI induced cognitive deficits ^16,19^, we measured learning and memory function one week, one month and eight months after TBI using the radial arm water maze (Figure 1G). As previously reported ^38,39^, mice were first trained to locate a platform hidden under opaque water placed in one of the eight arms using navigational cues placed in the room over one or two learning days of 9 trials or 6 trials, respectively (see Methods). Spatial memory was then tested by running a memory probe (3 consecutive trials) twenty-four hours and one week after training. The number of non-target arm entries before locating the escape platform (“errors”) were used as a measure of learning and memory deficits. Errors from three consequent trials were averaged in blocks. Of note, no sex differences were observed in the RAWM performance of female and male mice (Supplementary Figure 2) and were therefore plotted together. In line with previous studies, TBI induced a significant impairment in learning and memory as early as one week after injury (Two-way RM ANOVA, TBI effect: *F*1,30 = 9.783, *p* = 0.0039; time effect: *F*2.654,71.66 = 23.33, *p* < 0.0001) compared to sham mice (Figure 1H. Cognitive performance continued to be affected in injured mice at one month after TBI (Two-way RM ANOVA, TBI effect: *F*1,28 = 9.803, *p* = 0.0041; time effect: *F*3.120,87.35 = 23.41, *p* < 0.0001) (Figure 1I). Remarkably, cognitive deficits persisted in injured Ccr2-creER^T2^::Ai14D mice for up to eight months after trauma (Two-way RM ANOVA, TBI effect: *F*1,32 = 5.792, *p* = 0.0220; time effect: *F*3.385,108.3 = 27.18, *p* < 0.0001) (Figure 1J). These results demonstrate that Ccr2-creER^T2^::Ai14D female and male mice develop early and long-lasting deficits in spatial learning and memory after injury. Notably, we demonstrate for the first time that CCI-induced cognitive deficits as measured by RAWM persist for up to 8 months in injured mice.

### 3.3 MDMs accumulate in the pericontusional region and undergo phenotypic transition

To map the spatial localization of MDMs in the injured brain, we acquired z-stacked tiled images of the pericontusional area at 3 different coordinates from rostral to caudal (Figure 2A). As shown in the representative images (Figure 2B), most of the tdTomato+ MDMs accumulate in the cavitation area, few of them were found lining in the third ventricles and choroid plexus and in the CA1/DG regions of the hippocampus at 7 days. At chronic time points, cells were mainly localized in the thalamus under the ipsilateral hippocampus and in the choroid plexus (Figure 2B). No tdTomato+ cells were found in the contralateral hemisphere (data not shown). In line with the cell numbers observed by flow cytometry, tdTomato+ infiltrated macrophages at 1 and 8 months after injury were less numerous (Figure 2B). To determine if infiltrated tdTomato+ cells were differentiating into microglia, we analyzed the colocalization of tdTomato+ cells with ionized calcium-binding adapter molecule1 (Iba1) (expressed by macrophages) and the purinergic receptor P2ry12 (a microglia specific marker) across the pericontusional region (Figure 2C). We found a significant increase in the % of tdTomato+ cells that expressed Iba1 over time after engraftment (One-way ANOVA, *F* = 51.17, *p* < 0.0001, Figure 2D, Supplementary Figure 3). Notably, no tdTomato+ Iba1+ cells expressed P2ry12 at 7 days, but we observed a trend towards increased P2ry12 expression, though not statistically significant, at 8 months compared to 30 days (Figure 2E, Supplementary Figure 3). Next, we analyzed morphological changes in cell volume across the 3 selected hippocampal regions (anterior, central and posterior, Figure 2A). We observed a general reduction in cell volume from 7d to 30d and 8mos in tdTomato+ cells that also express Iba1+ (One-way ANOVA, Anterior *F* = 270.7, *p* < 0.0001, Central *F* = 184.9, *p* < 0.0001, Posterior *F* = 322.6, *p* < 0.0001) (Figure 2F). Interestingly, tdTomato+Iba1+P2ry12+ cells’ volume increased from 1 to 8 months after engraftment (Figure 2G). No tdTomato+ cells were found in coronal sections of sham animals at any time points (Supplementary Figure 4). These results show that MDMs preferentially accumulate in the injured region, undergo morphological changes and acquire the phagocytic marker Iba1 over time after engraftment. However, they do not fully differentiate into microglia.

**Figure 2.**
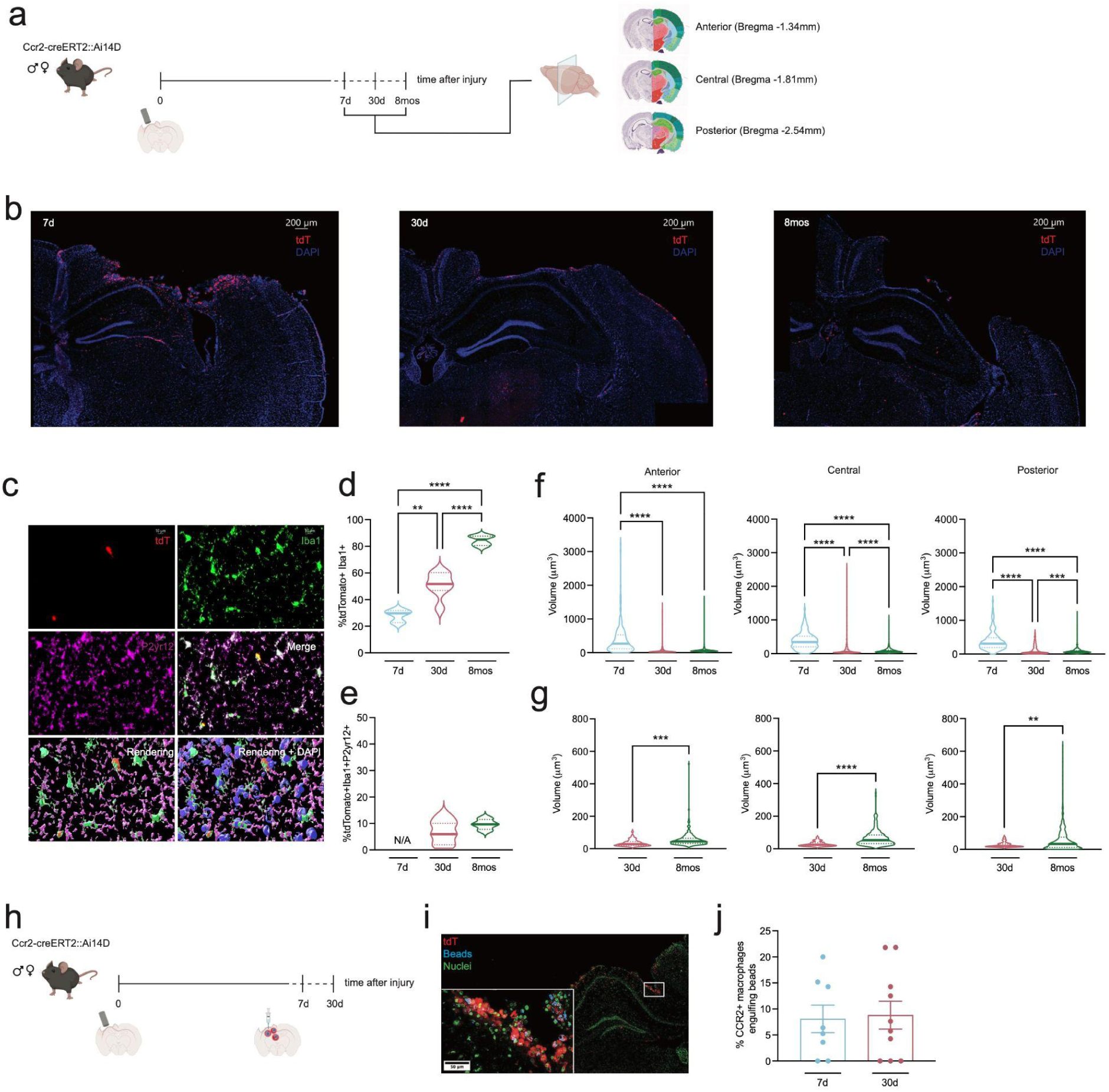
MDMs infiltrate the pericontusional region and maintain their phagocytic ability after engraftment. (a) Experimental design - Ccr2-creER^T2^::Ai14D mice were injured using the controlled cortical impact TBI model and brain samples were collected at different time points. For labeled macrophages visualization, pericontusional regions (top quarter of a coronal brain section) were acquired as 20x z-stacked tiled images. For each mouse, cavitation images were acquired at 3 different coordinates from Bregma: -1.34mm (“Anterior”), -1.81mm (“Central”) and -2.54mm (“Posterior”). Anatomical annotations from the Allen Reference Atlas – Mouse Brain, https://atlas.brain-map.org/ (b) Representative images of pericontusional regions of Ccr2-creER^T2^::Ai14D mice at different time points. (c) Representative example of machine learning-based surface rendering (Imaris 10.2.0) (d) % of tdTomato+ cells that express Iba1+ (e) Volume (µm^3^) of tdTomato+Iba1+ cells across the 3 selected coronal sections. (f) % of tdTomato+ cells that express Iba1 and P2yr12. (g) Volume (µm^3^) of tdTomato+Iba1+P2yr12+ cells across the 3 selected coronal sections. Violin plots depict the distribution of samples collected from *n* = 3/7 mice/group. \*\**p*<0.01, \*\*\*\**p*<0.0001 (One-way ANOVA and Unpaired *t*-test). (h) Experimental design - Ccr2-creER^T2^::Ai14D were injured using the controlled cortical impact TBI model and phagocytosis capacity was measured at 7 and 30 days after injury. Fluorescent beads (2μm diameter) were injected in the ipsilateral hippocampus, and brains were collected 3 days after for IHC analysis. (i) Representative confocal image of infiltrated macrophages (tdTomato, red) engulfing injected beads (blue) in the pericontusional area. Nuclei are visualized in green. (j) % of tdTomato+ macrophages that engulfed one or more fluorescent beads was quantified for each acquired image. Each dot is the average of 2/4 images acquired for each animal. Bars depict mean ± SEM of *n* = 8/10 mice/group. Female samples are shown as triangles and males as circles. \**p*<0.05, \*\**p*<0.01, \*\*\**p*<0.001 (Unpaired *t*-test).

### 3.4 MDMs maintain their phagocyte competency after engrafting the injured brain

Macrophages are professional phagocytes. To test the phagocytic capacity of TBI-induced MDMs and examine how this function changes over time after infiltration, we used our well established *in vivo* phagocytosis assay^38,59^. Blue fluorescent beads (2μm diameter) were injected in the ipsilateral hippocampi of injured Ccr2-creER^T2^::Ai14D mice on day 4 and 27 post-injury. Three days later (7 dpi or 30 dpi), brains were harvested for IHC analysis (Figure 2H). We quantified the percentage of tdTomato+ macrophages that engulfed one or more fluorescent beads on 40x Z-stack tiled images of the top half quarter of the mouse brain coronal section (pericontusional region) (Figure 2I). Strikingly, macrophages’ capability to phagocyte synthetic beads was unchanged between 7 and 30 dpi (Figure 2L). Of note, no sex differences were observed (Supplementary Figure 5). These data are the first to demonstrate that MDMs that engraft the brain after TBI are functional and can phagocyte, both at subacute and chronic time points.

### 3.5 Long after brain engraftment MDMs show a transcriptomic identity distinct from microglia that overlaps with established signatures of aging, senescence, and disease

After establishing long-term presence of MDMs in the injured brain, we investigated how the transcriptomes of these cells change over time after injury. MDMs (tdTomato+) were sorted out of the CD11b+CD45+ cell population in the whole Ccr2-creER^T2^::Ai14D brain at 7 days, 30 days, and 8 months after TBI for bulk RNA sequencing. Microglia was sorted as CD11b+, CD45^mid/low^, tdTomato- at 8 months after injury (Figure 3A). To characterize the molecular phenotypes of each group, we scored each sample by its enrichment of Reactome pathways^60^ using the singscore package^47^. We performed sparse partial least squares linear-discriminant analysis (sPLS-DA) of the Reactome scores to identify biological pathways that distinguish each group. Visualization of the discriminant components revealed a temporal trajectory of MDMs, with the first component capturing their dynamic differentiation over time following brain engraftment. The second component highlighted a distinct separation between MDMs and microglia at 8 months post-TBI, while the third component demonstrated a separation between MDMs at 7 and 30 days after injury (Figures 3B, Supplementary Figure 6). These results suggest that MDMs collected at 8 months after infiltration exhibit an intermediate transcriptional profile between MDMs at early time points and microglia. Pathways that were upregulated in the MDMs at 7 and 30 days relative to the 8 month samples included histone modifications and DNA damage (Figure 3C). Early in the TBI injury response, MDMs at 7 days were enriched for immune signaling and responses to cell stress and transitioned to responses to neuronal signaling and maresin signaling at 30 days. At 8 months after injury, MDMs upregulated terms related to immune system activation (e.g. Complement cascade, Interleukin-2 signaling, Synthesis of Leukotrienes (LT) and Eoxins(EX)), cell-cell interactions (e.g. Regulation of CDH11 Expression and Function, Regulation of Homotypic Cell-Cell Adhesion, Signaling to RAS), and signal transduction (e.g. Cell death signaling via NRAGE, NRIF and NADE) relative to resident microglia, which were enriched for terms related to cell maintenance (Receptor Mediated Autophagy and Cilium Assembly) (Figure 3C). We observed similar separation between groups when we performed the same analysis with the ImmunoSigDB^61^ dataset, a compendium of 5000 gene-sets from multiple immunology studies (Supplementary Figure 6). Heatmaps and volcano plots of differentially expressed genes between MDMs and microglia at 8 months after injury showed that while macrophages express higher levels of *Ccl2*, *Ccr2*, *Lyz2* and F4/80 (*Adgre1*), signature genes such as *Sall1, Hexb, Tmem119* and *P2ry12* were instead only upregulated in microglia (Figure 3D and 3E). These data demonstrate that MDMs retain their distinct transcriptomic profile even long after brain engraftment.

**Figure 3.**
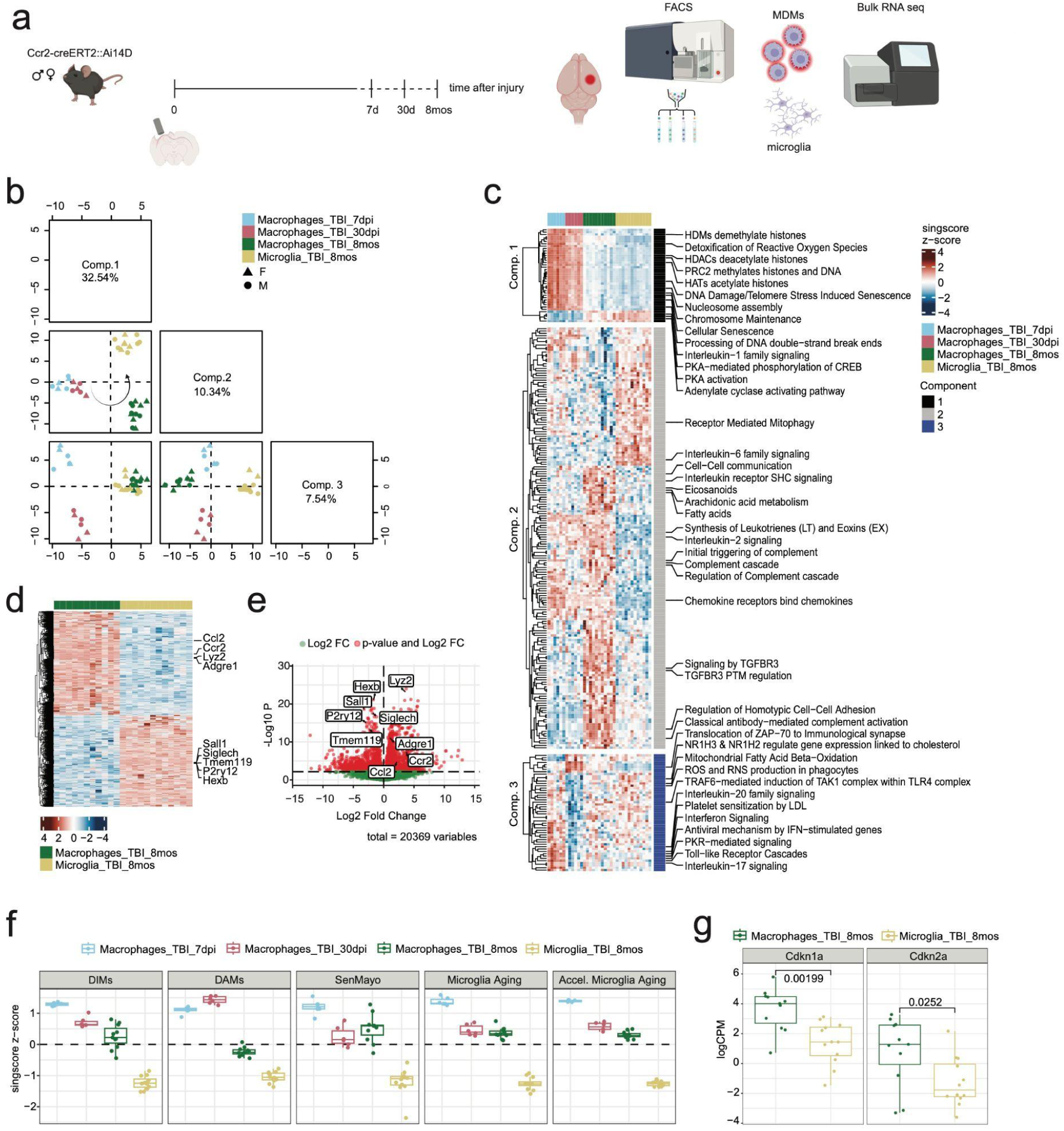
Transcriptomic signatures of MDMs after TBI. (a) Experimental design - Ccr2-creER^T2^::Ai14D mice were injured using the controlled cortical impact TBI model and brain samples were collected at different time points. Microglia (tdTomato-) and MDMs (tdTomato+) cells were isolated from the CD11b+CD45+ brain cell population by FACS and processed for bulk RNA sequencing. (b) Projections of samples onto the first three components of sPLS-DA analysis of Reactome singscore scores from TBI macrophages at 7dpi, 30dpi, and 8mos, with 8mos TBI microglia. (c) Heatmap of Reactome singscore z-scores for significant terms for each component selected by sPLS-DA. The top terms for discriminating each group are shown. (d) Heatmap and (e) volcano plot of DEGs from TBI macrophages and microglia 8 months after injury. (f) Reactome singscore z-scores for reference signatures of disease-inflammatory macrophages (DIMs), disease-associated microglia (DAMs), SenMayo, Microglia Aging, and Accelerated Microglia Aging (see methods). (g) Expression (logCPM) of *Cdkn1a (p21)* and *Cdkn2a (p16)* in TBI macrophages and microglia 8 months after injury (Unpaired *t*-test, *p-value* in the graph). Animals (6-12 for each condition) are individually plotted. Box plots depict data quartiles.

Next, we compared the MDMs transcriptomes to recently published well-established macrophages/microglia cell signatures. We focused on the “Disease Inflammatory Macrophages” (DIM) population, which accumulates in the mouse brain with aging and in neurodegenerative conditions ^62^, and with the “Disease Associated Microglia” (DAM) signature^63^. As shown in Figure 3F, both DIM and DAM signatures were enriched in MDMs, especially at the early time points after TBI (Figure 3F). This indicates that MDMs display a unique signature associated with disease that does not occur in microglia. Next, we compared our data with the SenMayo gene set, a senescence list of 125 genes validated in aged human and mouse samples ^64^ and found that MDMs are enriched in terms related to senescence compared to microglia (Figure 3F). Similar results were observed when comparing our infiltrated macrophages with microglia isolated from 24 months old mice (“Microglia Aging”) and from Ercc1Δ/KO mice that display features of accelerated aging (“Accel. Microglia Aging”)^65^(Figure 3F). Finally, we assessed the gene expression of *Cdkn1a (p21)* and *Cdkn2a (p16)*, two of the most widely used genetic markers of senescence, not included in the SenMayo signature ^64,66^. As shown in Figure 3G, mRNA levels of *p21* and *p16* were significantly higher in MDMs at 8 months after TBI when compared to microglia. These results reveal that MDMs transcriptomic signature is associated with aging, senescence and disease that is distinct from microglia and further support the causative role for the development of cognitive deficits we previously reported.

### 3.6 MDMs share a common core transcriptomic signature across fate mapping models and brain perturbations

We validated our transcriptomic findings using Ms4a3-cre::Ai14D mice, a fate mapping model complementary to the Ccr2-creER^T2^ system^34^. In these mice, all the cells differentiated from granulocyte monocyte progenitor (GMP), including Ly6C^hi^ monocytes, are constitutively labeled ^35^ (Figure 4A). Of note, Ms4a3-cre::Ai14D mice displayed TBI-induced cognitive impairments in RAWM similar to Ccr2-creER^T2^::Ai14D mice (Supplementary Figure 7). MDMs (tdTomato+), microglia (CD11b+, CD45^mid/low^, tdTomato-) and “classical” monocytes (Ly6G-,Ly6C^hi^) were sorted from brain and blood of female and male Ms4a3-cre::Ai14D mice at 7 and 30 days post TBI (Figure 4A). Ms4a3+ MDMs displayed higher levels of *Ccr2* mRNA compared to microglia, comparable with what we observed in the Ccr2+ dataset (Figure 4B). Microglia specific genes were enriched in microglia vs Ms4a3+ MDMs at both time points, confirming that infiltrated macrophages retain their identity (Figure 4C-F). We then corroborated the transcriptomic signature associated with aging, senescence and disease in Ms4a3+ MDMs in comparison to sham microglia, with the exception of the DAM signature (Figure 4G).

**Figure 4.**
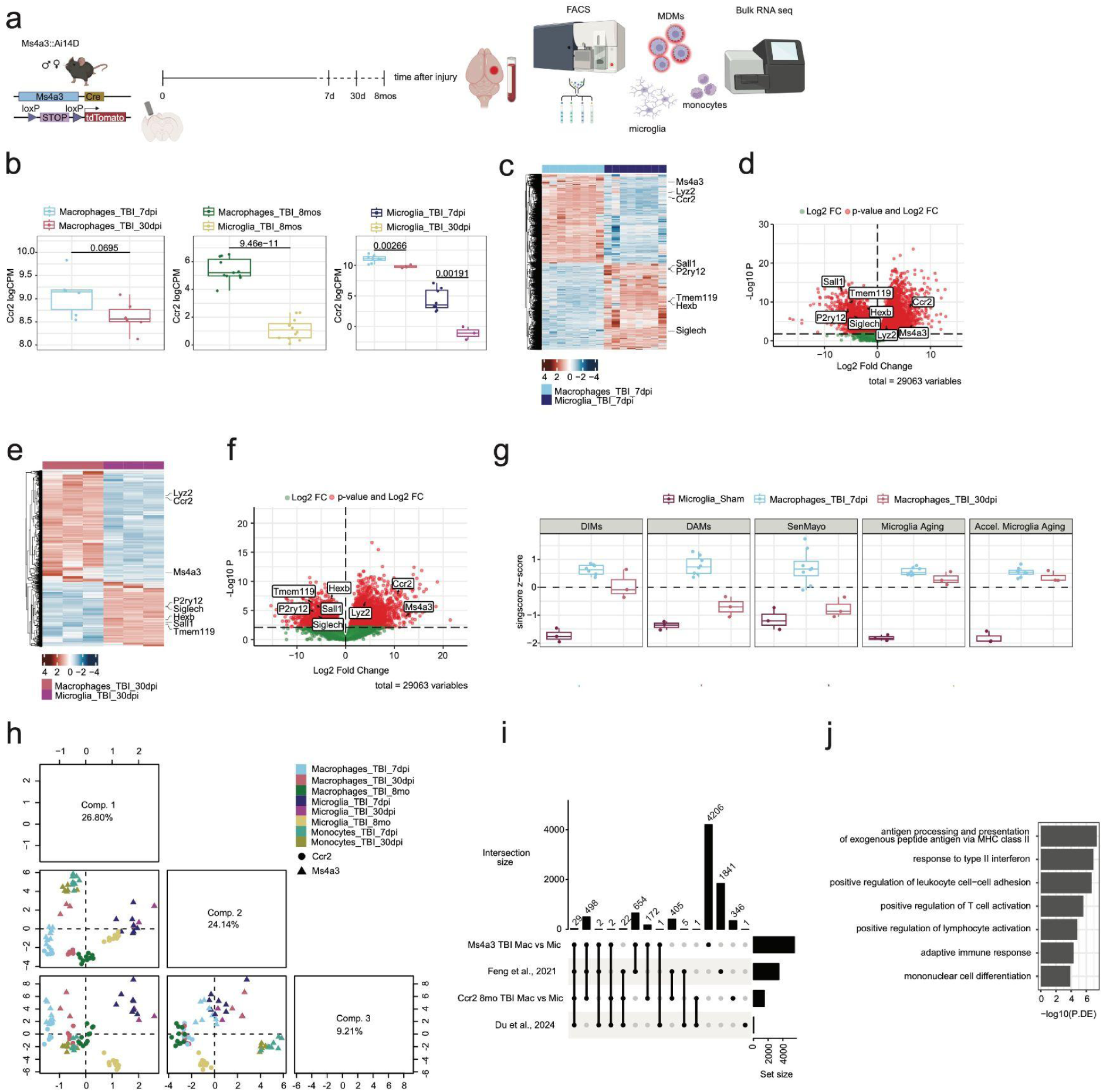
MDMs share a common core transcriptomic signature. (a) Experimental design - Ms4a3-cre::Ai14D mice were injured using the controlled cortical impact TBI model and brain samples were collected at different time points. MDMs (tdTomato+), microglia (CD11b+, CD45^mid/low^, tdTomato-) and inflammatory monocytes (Ly6G-,Ly6C^hi^) were sorted from brain and blood by FACS and processed for bulk RNA sequencing. (b) LogCPM of Ccr2 transcripts across datasets (Unpaired *t*-test, *p value* in the graph). (c) Heatmap of DEGs from TBI macrophages and microglia 7 days after injury. (d) Volcano plot of DEGs from TBI macrophages and microglia 7 days after injury. (e) Heatmap of DEGs from TBI macrophages and microglia 30 days after injury. (f) Volcano plot of DEGs from TBI macrophages and microglia 30 days after injury. (g) Singscore z-scores for reference datasets. (h) Projections of TBI samples from Ccr2-creER^T2^::Ai14D and Ms4a3-cre::Ai14D models onto the first three components of sPLS-DA analysis of singscore scores for cell type signatures from the Molecular Signatures Database (MSigDB). (i) UpSet plot of the overlaps between DEGs from different datasets (see Methods). (j) Selected GO:BP terms enriched in the list of the 29 genes commonly enriched in MDMs. Animals (3-12 for each condition) are individually plotted. Box plots depict data quartiles.

We next combined the transcriptomic data of microglia, monocytes, and MDMs from the Ccr2 and Ms4a3 datasets, and mapped them together onto cell type signatures from the Molecular Signatures Database (MSigDB)^67^ using singscore. sPLS-DA analysis of the resulting scores revealed general clustering by cell identity and separation by time point (Figure 4H, Supplementary Figure 8). Strikingly, these results confirm that MDMs from two different genetic models are transcriptomically distinct from microglia and monocytes, regardless of how long they have been engrafted in the brain. Lastly, we investigated if MDMs share a core genetic signature, across different models and experimental conditions. For this purpose, we compared our datasets with other bulk transcriptomic datasets of brain-engrafted macrophages available in the literature ^59,68^. We identified 29 genes that were commonly upregulated in MDMs (Supplementary Table) (Figure 4I). These genes were enriched for Gene Ontology Biological Processes related to immune response pathways such as antigen processing and presentation, immune cells activation, and differentiation (Figure 4L). These results strengthen the concept of long-lasting MDMs as distinct cell populations that share a core signature that is consistent across different models and experimental conditions.

## 4. Discussion

Using two independent and complementary fate mapping systems, we described for the first time a population of MDMs that persist in the mouse brain parenchyma for up to 8 months after injury, a time point when we still measure long-lasting cognitive deficits.

While we have previously demonstrated the functional role of MDMs in the development of cognitive deficits following TBI ^16,19^, it remained unknown the fate of MDMs upon engraftment. We and others have previously shown that the accumulation of these cells in the injured brain is temporally restricted ^69,70^, as these cells “disappear” from the injured brain after a few days^16^. The explanation behind this phenomenon lies in the use of Ccr2^RFP/+^ knock-in mice, where infiltrated macrophages will stop expressing Ccr2 and therefore their fluorescent tag, after *in situ* reprogramming^37^. Linear tracing using fate mapping models is a useful tool to unravel the long-term trajectories of MDMs^33^. By using this approach, we were able to unambiguously show the presence of labeled MDMs in the mouse brain for up to 8 months post TBI, challenging the assumption about their transient nature^69^. Of note, the number of infiltrated macrophages we detected subacutely (7dpi) in injured mice (around 6000/whole brain) was higher than what we published before in CX3CR1^+/GFP^Ccr2^+/RFP^ mice (less than 500/ipsilateral hemisphere at peak accumulation)^16^, and what Doran et al., found one week after TBI in WT mice (around 1000 cells/ipsilateral hemisphere)^71^. Such number disparity can be explained by the different mouse models deployed, and demonstrates the significant advantages of using a more rigorous fate mapping system. The number of brain infiltrated macrophages that persist in the brain after infiltration decreases from subacute time points (7 dpi, around 6000 cells) to chronic time points (30 dpi, around 2000 cells; 8 months post injury, around 600 cells). Leveraging existing TBI studies^69^, we hypothesize that a portion of infiltrated MDMs dies after the first wave of invasion, while a smaller niche (10%) durably persists in the injured brain parenchyma. Building on our previous work demonstrating that preventing MDMs infiltration into the brain can prevent TBI-induced cognitive deficits^16^, we theorize that the remaining MDMs contribute to the long-term cognitive impairments observed here for the first time at 8 months after TBI.

This durable population of MDMs tend to localize in the ipsilateral hippocampus, the thalamus underneath and the choroid plexus, differently from subacute time points when most of the infiltrated cells accumulate alongside the cavitation. We speculate that after having fulfilled their acute function of debris clearance and tissue remodeling around the injured area, engrafted macrophages integrate into the network of resident macrophages. Supporting this hypothesis, we observed a reduction in cell volume at chronic time points compared to 7 days post-injury. At 7 days, the larger cell volume likely reflects an enhanced phagocytic phenotype, corresponding to an increased demand for tissue clearance and repair subacutely after injury ^72^. Moreover, the % of tdTomato+ MDMs expressing the macrophages/microglia marker Iba1 increases over time after infiltration, reaching ∼85% at 8 months. However, only ∼10% of MDMs stained positive also for P2ry12, a well-established microglia specific marker^73^. This suggests that MDMs do not fully differentiate into resident microglia, even long after engraftment. Our results are consistent with other fate mapping studies showing that MDMs do not contribute to the resident microglia pool during brain infection^,74^ and after microglia depletion^,68^. Nevertheless, we show for the first time that MDMs are competent phagocytes both subacutely and chronically after TBI, in line with our previous findings in brain-engrafted macrophages (BEMs) that populate the brain after microglia depletion and have comparable phagocytic capacity^59^. An additional and conclusive proof that infiltrated macrophages maintain their distinct identity comes from the bulk RNA-Seq data. By using two distinct yet complementary fate mapping systems (Ccr2-creER^T2^ and Ms4a3-cre)^75^, we showed the MDMs’ transcriptomics undergoes dynamic changes after engraftment. Notably, even though MDMs 8 months after TBI exhibit an intermediate transcriptome between early time points macrophages and microglia, they still maintain their unique signature and don’t express microglia-specific genes. By comparing our data with reference datasets ^62,63,64,46^ we found that MDMs show a transcriptomic signature associated with disease, aging and senescence. We propose that this could contribute to the long-lasting cognitive deficits observed following TBI, as we have previously shown that preventing the entry of MDMs into the brain can avert the development of cognitive impairments ^16^. Finally, we generated a list of 29 core genes that are consistently shared across different published dataset focused on infiltrated macrophages ^59,68^ and have been previously implicated as drivers of critical processes including neuroinflammation, cellular damage, and repair mechanisms in TBI (e.g. *Cybb ^76^, Atp2b1 ^77^, Ifnar1 ^78^, Cd74 ^79^, Mrc1 ^80^, Tgfbi ^81^)*. Our findings not only will facilitate the identification of these cells in future transcriptomic studies but also provide valuable insights for developing targeted therapeutic strategies aimed at specific subpopulations of infiltrated macrophages.

## 5. Acknowledgments

We thank Professor Burkhard Becher (University of Zurich) for providing Ccr2-creER^T2^-mKate2 mice and Professor Florent Ginhoux (Singapore Immunology Network (SIgN), A*STAR) for providing Ms4a3-cre mice.

We thank the Genomics CoLabs of the University of San Francisco, California (UCSF) that run part of the Bulk RNA sequencing analysis.

We thank Eric Griffis (Altos Labs) for helping with the acquisition of images for the phagocytosis assay, Yuewen Zheng (Altos Labs) for her contribution to the sequencing analysis, Aniket Tolpadi (Altos Labs) for assisting with processing Imaris raw data, Shruti Suresh (Altos Labs), Ariana Wei (Altos Labs), Valentina Frattini (UCSF), Kim Chung (UCSF) for helping with sample processing.

Images were created with BioRender.com

## 6. Authors Contribution

Conceptualization: M.S.P., B.A.Y., S.R., X.F., K.K.

Experimental work: M.S.P., W.L., R.S.

Data analysis: M.S.P., B.A.Y., E.S.F., V.P., with the support from M.M., S.T.

Manuscript writing: M.S.P., S.R., B.A.Y.

All the authors made contributions to the manuscript editing.

## 7. Data and code availability

### Data

| Data | Citation | Source |
| --- | --- | --- |
| Bulk RNA-Seq of CD11b+ CD45+ cells from whole brains of Ccr2-creER<sup>T2</sup>::Ai14D | This paper | GEO: GSEXXXXX |
| Bulk RNA-Seq of CD11b+ CD45+ cells from whole brains of Ms4a3-cre::Ai14D | This paper | GEO: GSEXXXXX |
| Brain-engrafted macrophages | <sup>82</sup> | None |

