## Supplementary Figures for "Fate mapping of peripherally derived macrophages reveals a long-lasting engrafted population that maintains a distinct transcriptomic profile for up to 8 months after Traumatic Brain Injury"

### Supplementary Figure 1 - Blood labeling at the time of injury

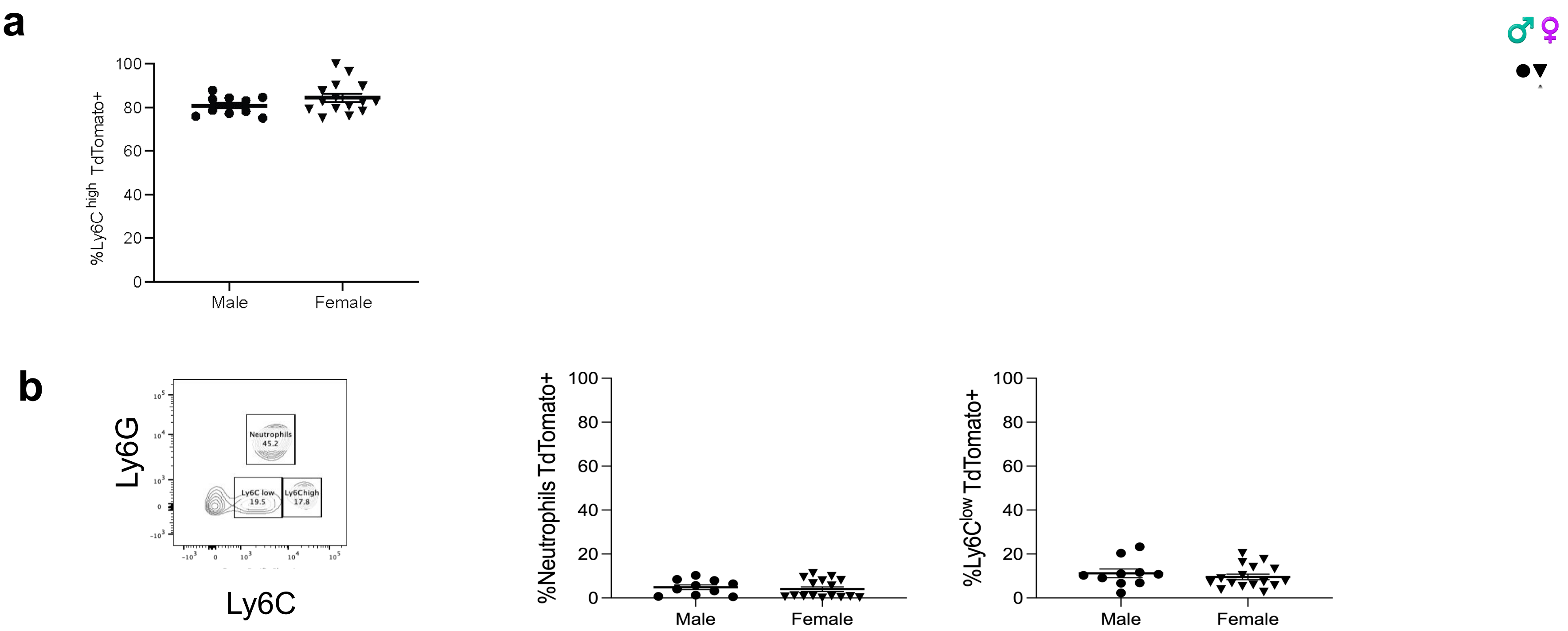


**Supplementary Figure 1 -**
(**a**) Off-target labeling of neutrophils (gated as CD11b+Ly6C+Ly6G+) and patrolling monocytes (gated as CD11b+Ly6G-Ly6Clow) after 3 doses of tamoxifen (at the time of injury) was below 15%. (**b**) Labeling efficiency expressed as percentage of blood Ly6C$^{high}$ monocytes that were tdTomato+ at the day of injury (3 doses of tamoxifen, day 0). No sex differences were observed. Individual animals are plotted. Bars are the mean of the examined variable ± SEM of *n* = 10/16 mice/group. Unpaired *t*-test.

### Supplementary Figure 2 - No sex differences in cognitive deficits after TBI

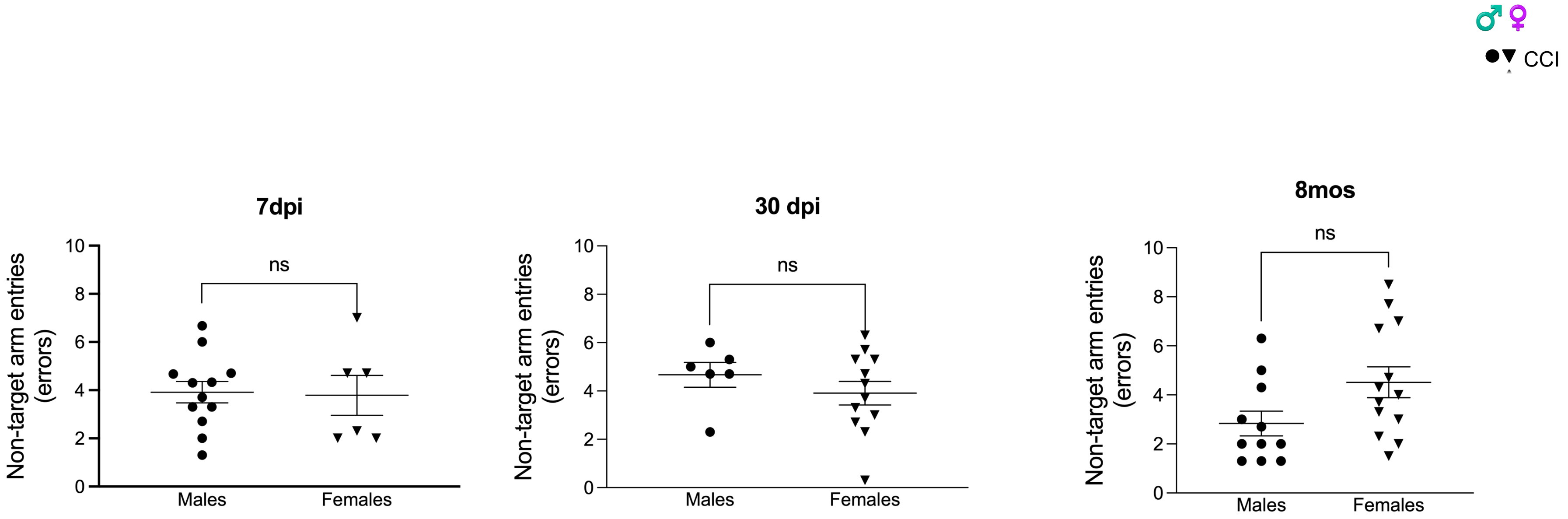


**Supplementary Figure 2 -**
Ccr2-creER$^{T2}$::Ai14D mice develop memory deficits after focal injury (CCI). No sex differences were observed in the memory probe performance of CCI mice. Male mice are coded in circles and female in triangles. Individual animals are plotted. Bars are the mean of the examined variable ± SEM of *n* = 6/13 mice/group. Unpaired *t*-test.

### Supplementary Figure 3 - tdTomato+ colocalization with Iba1 and P2yr12 - quantification by coordinates

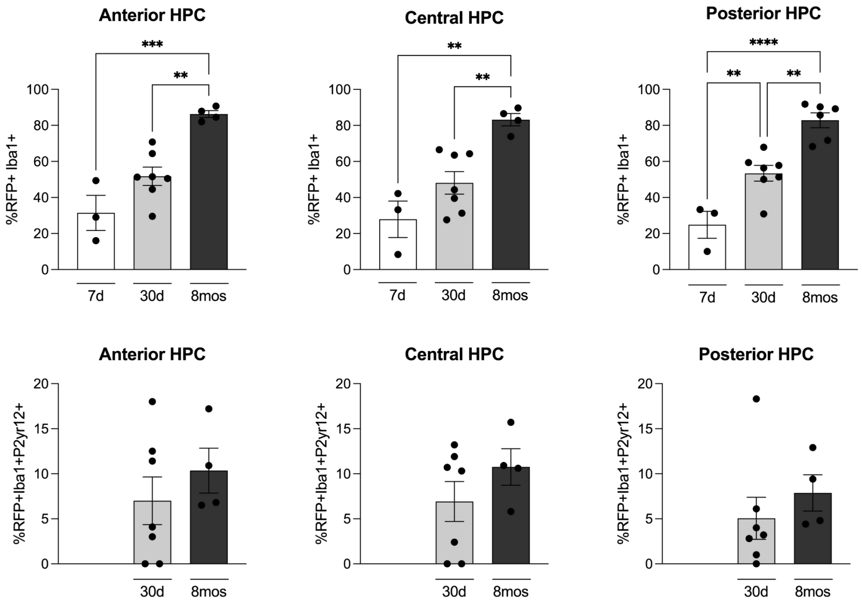


**Supplementary Figure 3 -**

For labeled macrophages visualization, pericontusional regions (top quarter of a coronal brain section) were acquired as 20x z-stacked tiled images. For each mouse, cavitation images were acquired at 3 different coordinates from Bregma: -1.34mm (“Anterior”), -1.81mm (“Central”) and -2.54mm (“Posterior”). Top: % of tdTomato+ cells that express Iba1+. Bottom: % of tdTomato+ cells that express Iba1 and P2yr12. Samples (*n* = 3/7 mice/group) are individually plotted. One-way ANOVA with Tuckey multiple comparisons test.

### Supplementary Figure 4 -
### tdTomato+ cells don’t infiltrate the brain parenchyma of Sham Ccr2-creER$^{T2}$::Ai14D mice

**7d** **30d** **8mos**

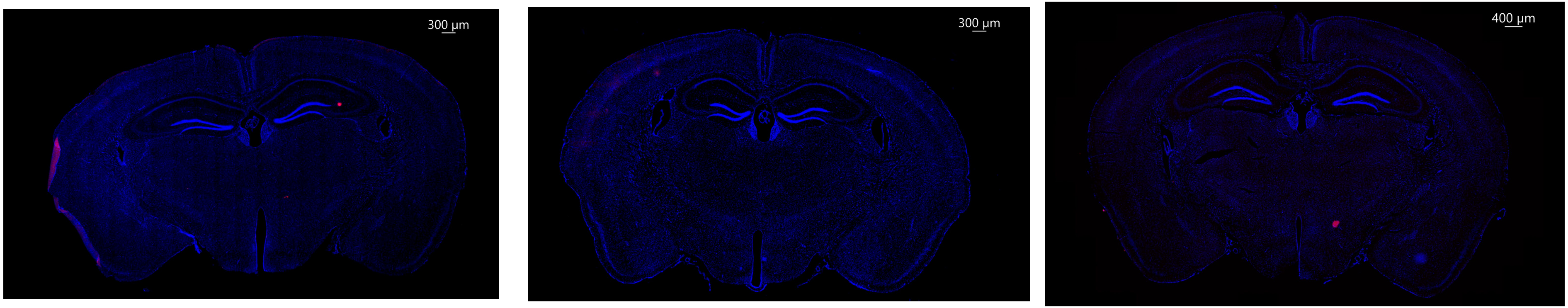


**Supplementary Figure 4 -**
No tdTomato+ cells were found in the coronal sections of sham Ccr2-creER$^{T2}$::Ai14D mice

### Supplementary Figure 5 - No sex differences in phagocytosis

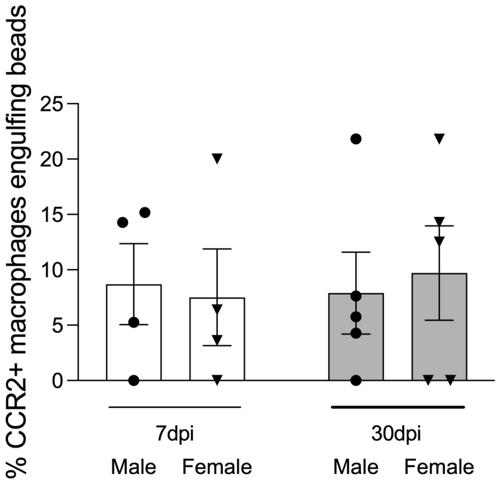


| | | | | | |
|---|---|---|---|---|---|
| ANOVA table | SS (Type III) | DF | MS | F (DFn, DFd) | P value |
| Interaction | 9.977 | 1 | 9.977 | F (1, 14) = 0.1361 | P=0.7178 |
| Sex | 0.4438 | 1 | 0.4438 | F (1, 14) = 0.006052 | P=0.9391 |
| Time after injury | 2.293 | 1 | 2.293 | F (1, 14) = 0.03127 | P=0.8622 |
| Residual | 1027 | 14 | 73.33 | | |

**Supplementary Figure 5 -**
No sex differences were observed in the phagocytosis capability of TBI-induced infiltrated macrophage, as shown by the Two Way ANOVA analysis.

### Supplementary Figure 6 - Infiltrated macrophages (Ccr2) transcriptomes mapped onto ImmuneSigDB dataset

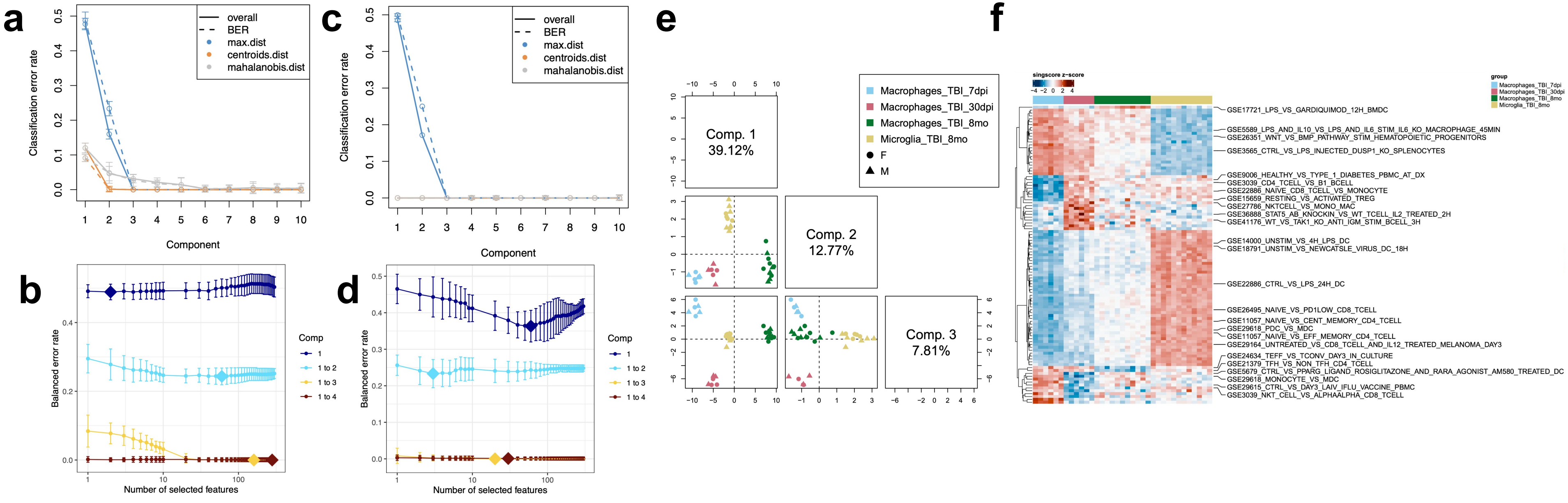


**Supplementary Figure 6 -** Classification error rates per number of sPLS-DA components for the (**a**) Reactome and (**c**) ImmuneSigDB databases. 3 components were selected for the final models for both databases. Balanced error rate per number of selected features per discriminant component for the (**b**) Reactome and (**d**) ImmuneSigDB databases. (**e**) Projections of samples onto the first three components of sPLS-DA analysis of ImmuneSigDB singscore scores from TBI macrophages at 7dpi, 30dpi, and 8mo, with 8mo TBI microglia. (**f**) Heatmap of ImmuneSigDB singscore z-scores for significant terms for each component selected by sPLS-DA. The top terms for discriminating each group are shown.

### Supplementary Figure 7 - Ms4a3::Ai14dT RAWM

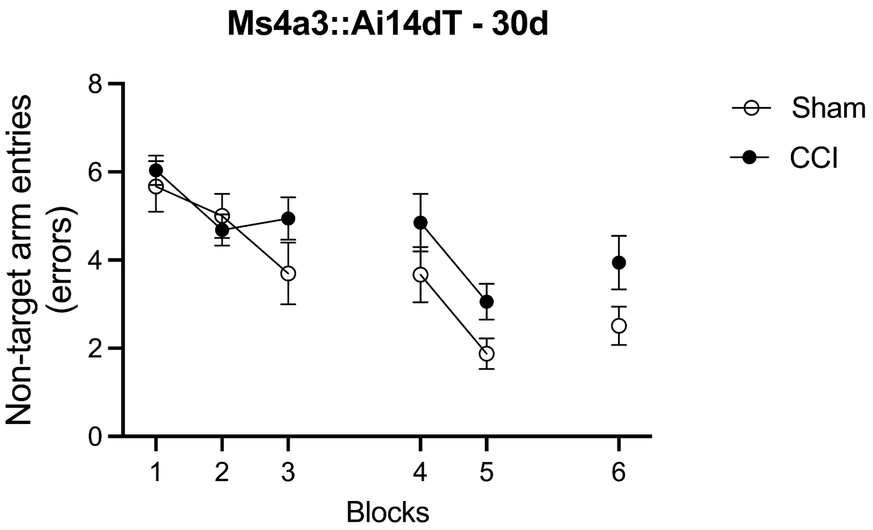


| | | | | | |
|---|---|---|---|---|---|
| Source of Variation | % of total variation | P value | P value summary | Significant? | Geisser-Greenhouse's epsilon |
| Time x TBI | 1.805 | 0.4649 | ns | No | |
| Time | 22.28 | <0.0001 | **** | Yes | 0.7333 |
| TBI | 3.395 | 0.0444 | * | Yes | |
| Subject | 20.61 | 0.0065 | ** | Yes | |

**Supplementary Figure 7 -**
Ms4a3::Ai14dT mice show learning and memory deficits in the RAWM 30 days after TBI. Data are expressed as mean of the examined variable ± SEM of *n* = 11/18 mice/group. **p*<0.05, ***p*<0.01, *****p*<0.001 (Two way RM ANOVA).

### Supplementary Figure 8 - Infiltrated macrophages transcriptomes (Ccr2 and Ms4a3 combined) after brain engraftment mapped onto the Molecular Signatures and Reactome database

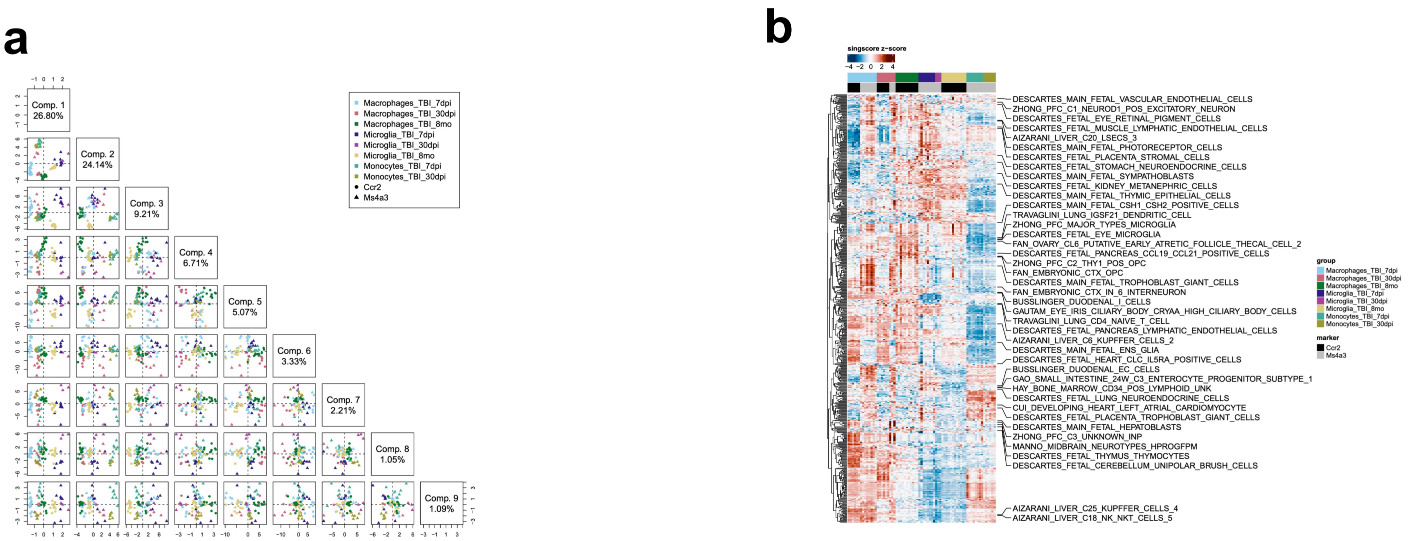


**Supplementary Figure 8 -** (**a**) Projections of samples onto the first 10 components of sPLS-DA analysis of MSigDB cell type signature singscore scores from TBI samples. (**b**) Heatmap of cell type signature singscore z-scores for significant terms for each component selected by sPLS-DA. The top terms for discriminating each group are shown. (**c**) Classification error rates per number of sPLS-DA components. 10 components were selected for the final models for both databases. (**d**) Balanced error rate per number of selected features per discriminant component.

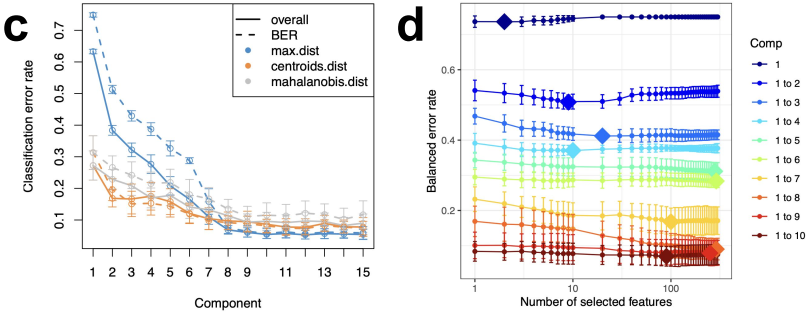
